# Nutrition of honeybees is constrained by the ratios of essential amino acids in pollen protein

**DOI:** 10.1101/2025.09.24.674040

**Authors:** Daniel Stabler, Jennifer A. Chennells, Eileen F. Power, Jennifer Scott, Giorgia Perri, Philip K. Donkersley, Sharoni Shafir, Geraldine A. Wright

**Affiliations:** School of Biological Sciences, Faculty of Environmental and Life Sciences, University of Southampton, University Road, Southampton, SO17 1BJ, UK; Department of Biology, University of Oxford, Oxford, OX1 3SZ, UK; Centre for Behaviour and Evolution, Institute of Neuroscience, Newcastle University, Newcastle upon Tyne, NE1 7RU, UK; Department of Nutrition, Food and Exercise Sciences, School of Biosciences, Faculty of Health and Medical Sciences. University of Surrey, Guildford, Surrey, GU2 7XH, UK; Lancaster Environment Centre, Lancaster University, Lancaster LA1 4YQ, UK; B. Triwaks Bee Research Center, Department of Entomology, Institute of Environmental Sciences, Robert H. Smith Faculty of Agriculture, Food and Environment, The Hebrew University of Jerusalem, Rehovot, 7610001, ISRAEL

## Abstract

Bees pollinate most of the world’s flowering plants and collect and eat floral pollen as their sole source of dietary protein, yet we know relatively little about nutritional constraints imposed on them by pollinivory. Pollen protein provides bees with essential amino acid (EAA) profiles that vary as a function of plant species, but the extent of this variation and its impact on bee feeding behaviour and performance are unknown. Here, we measured the EAA profiles of pollen, bee bread, honeybees, and royal jelly to understand natural variation in protein quality and tested how mismatches in EAAs relative to bee tissues impacted protein-to-carbohydrate regulation in adult workers. Bees fed diets with an EAA profile that matched their own tissues consumed more food, gained more weight, and ate proportionally more protein relative to carbohydrate (1:72 EAA:C) while those fed with pollen sources including bee bread ate proportionally less protein and less food overall. Deficiencies found in pollen led us to discover that nutrient balancing for protein and carbohydrate in bees was driven by the inverse relationship between quantities of the branched-chain amino acids relative to histidine in dietary protein. We predict that the creation of bee bread, a mixture of bee-collected pollen, is a general adaptation to pollen feeding that reduces the impact of imbalances in the EAA profile of pollen protein.

## Introduction

Bees are a unique group of herbivorous animals that feed almost exclusively on floral pollen and nectar. Their pollen-feeding habit is derived from ancestors that originally fed on insect prey (Murray et al., 2018). Bees co-evolved with flowering plants, and as plants diversified, they adapted to derive all the essential protein, lipids, and sterols they require to rear offspring from floral pollen. The nutritional value of pollen, however, varies considerably as a function of plant species, particularly in the composition of protein and lipid (Manning, 2001; Somerville and Nicol, 2006; Arien et al., 2015; Dufour et al., 2020; Vaudo et al., 2020; Jeannerod et al., 2022). Although pollen is often assumed to be a near perfect food for bees, it is the male gamete of plants and, unlike nectar, it is rarely produced solely as a reward for pollinators. Additionally, nectar composition is mainly carbohydrates and water (Nicolson, 2022), whereas pollen is composed of protein, fats, sterols, and many other compounds (e.g. polyphenols). For this reason, pollen is more metabolically expensive for a plant to produce; because it is the male gamete, its loss also directly impacts plant fitness. Given the conflict of interest between plant and pollinator, it is unlikely that all bee species have succeeded in shaping the nutritional value of pollen to suit their needs (Roulston and Cane, 2000). However, some obligate insect-pollinated species do produce pollen with higher protein contents than self- or wind-pollinated species (Hanley et al., 2008). There is some evidence that nutrient composition of pollen affects honeybee foraging efforts (Cook et al., 2003; Hendriksma et al., 2014; Vaudo et al., 2016; Ghosh et al., 2020; Yokota et al., 2024), but whether the differences in protein quality in pollen matter to nurse-aged honeybees that consume the pollen collected by foragers has never been explicitly tested.

Dietary protein is most animals’ main source of essential amino acids (EAAs): compounds that they must acquire for growth, maintenance, and reproduction. Protein from animal tissue is often a close match to what is needed by other animals, whereas protein from plant sources is much more variable in the profile of EAAs (MacArthur et al., 2021; Dai et al., 2022). Poor quality proteins can have a large impact on animal health and fitness if necessary EAAs are missing or too concentrated (Dai et al., 2022). For example, many plant proteins have relatively low quantities of lysine (Leinonen et al., 2018) and/or the branched chain amino acids (BCAAs) (isoleucine, leucine and valine) relative to animal tissue (MacArthur et al., 2021). Exclusively eating certain plant proteins can lead to deficiencies which cause developmental, immunological, and metabolic problems in mammals (Balnave and Brake 2002; MacArthur et al., 2021; Dai et al., 2022). Likewise, mismatches in the EAAs present in foods and those required by insects and other arthropods can affect development, survival, and fecundity (Piper et al., 2017; Guisande et al., 1999; Ma et al., 2022).

The EAA profile of pollen varies among plant species (Hsu et al., 2021). Though some studies have compared the EAA profile of pollen to the EAA recommendations from the studies of de Groot (1953) for honeybees (Weiner et al., 2010; Dufour et al., 2020), none have direct experimental evidence to show how such differences in EAA composition impact honeybee performance or behaviour. Some have shown the importance of honeybee preference for individual amino acids in foraging decisions (Hendriksma, et al., 2014), but largely ecologists have struggled to define the nutritional value of plant species’ pollen for honeybees, because no framework other than that of de Groot’s (1953) exists. Such information is critically important for the remediation of landscapes to improve habitat for honeybees and other pollinators (Baude et al., 2016).

Here, we studied how the EAA profiles of pollen influence macronutrient balancing of adult worker honeybees using the Geometric Framework for Nutrition (Raubenheimer and Simpson, 1999). We first identified the EAA profile of honeybees calculated from its exome. We also measured EAAs rendered after the protein hydrolysis of pupal tissues and of the pollen of a subset of the major families of flowering plants in the UK. Using these data, we compared honeybee performance on diets made of free EAAs designed to test how pollen’s EAA profiles affected the feeding behaviour and weight (a proxy for muscle tissues) of adult worker honeybees. In our final experiments, we explicitly tested how the ratios of specific EAAs in diet (e.g. leucine and histidine) impacted protein-to-carbohydrate balancing and the regulation of feeding. These experiments allowed us to pinpoint the EAAs that impact protein-to-carbohydrate regulation in food and to identify which plant families have the most suitable pollen for honeybees.

## Results

### A mismatch exists between bee EAA and pollen EAA

We measured the amino acids found in bee tissue (exome and pupal hydrolysis), royal jelly, bee bread (stored pollen), and hand-collected pollen from five different plant families (Brassicaceae, Fabaceae, Lamiaceae, Rosaceae, and Solanaceae). We also included the proportions of EAAs predicted by de Groot (1953). Using a canonical discriminant analysis (CDA), we tested whether it was possible to separate bee tissues from bee bread and hand-collected pollen using the proportions of the EAAs found in protein. Bee tissues differed most greatly from the EAAs found in pollen or bee bread (Figure 1A, Table S1, CDA, function 1, 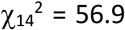 P < 0.001). Bee tissues, like other animal proteins, had a strong positive correlation in the presence of the branched chain amino acids (BCAAs: ile, leu, val) (Figure 1A, table S1, note: tryptophan not included), whereas pollen proteins had stronger correlations in the presence of arginine and histidine. Curiously, the EAA profile of bee bread was significantly different to both hand-collected pollen and to bee tissue (Table S1, CDA, function 2, 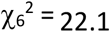, P = 0.001). Threonine was highest in bee bread (30% of total), and methionine and lysine were proportionally lower in this treatment than in any other (Figure 1A, table S1). As there was a clear mismatch between the EAAs found in bee tissues, bee bread, and pollen, we predicted that the EAA profiles could affect macronutrient balancing, and hence, the performance of bees.

### Experiment 1: Nutrient balancing for EAA and carbohydrates depends on the EAA profile in diet

For 10 days, we fed cohorts of newly emerged worker honeybees with one of nine diets (Table S3). These included an EAA profile that matched honeybees or bee products (Exome, Royal Jelly, or Bee Bread), the average EAA profile of the plant families in Figure 1A, or a diet with the EAAs present in equal quantities (Equal EAA). The measured values of honeybee pupae, royal jelly, and five plant families are presented in Table S2 and values were translated to proportions as reported in Table S3. The bees were given a choice of the treatment diet and a diet composed of sucrose with a lipid emulsion, so that they could balance their intake of carbohydrates independently of the treatment diet.

**Figure 1.**
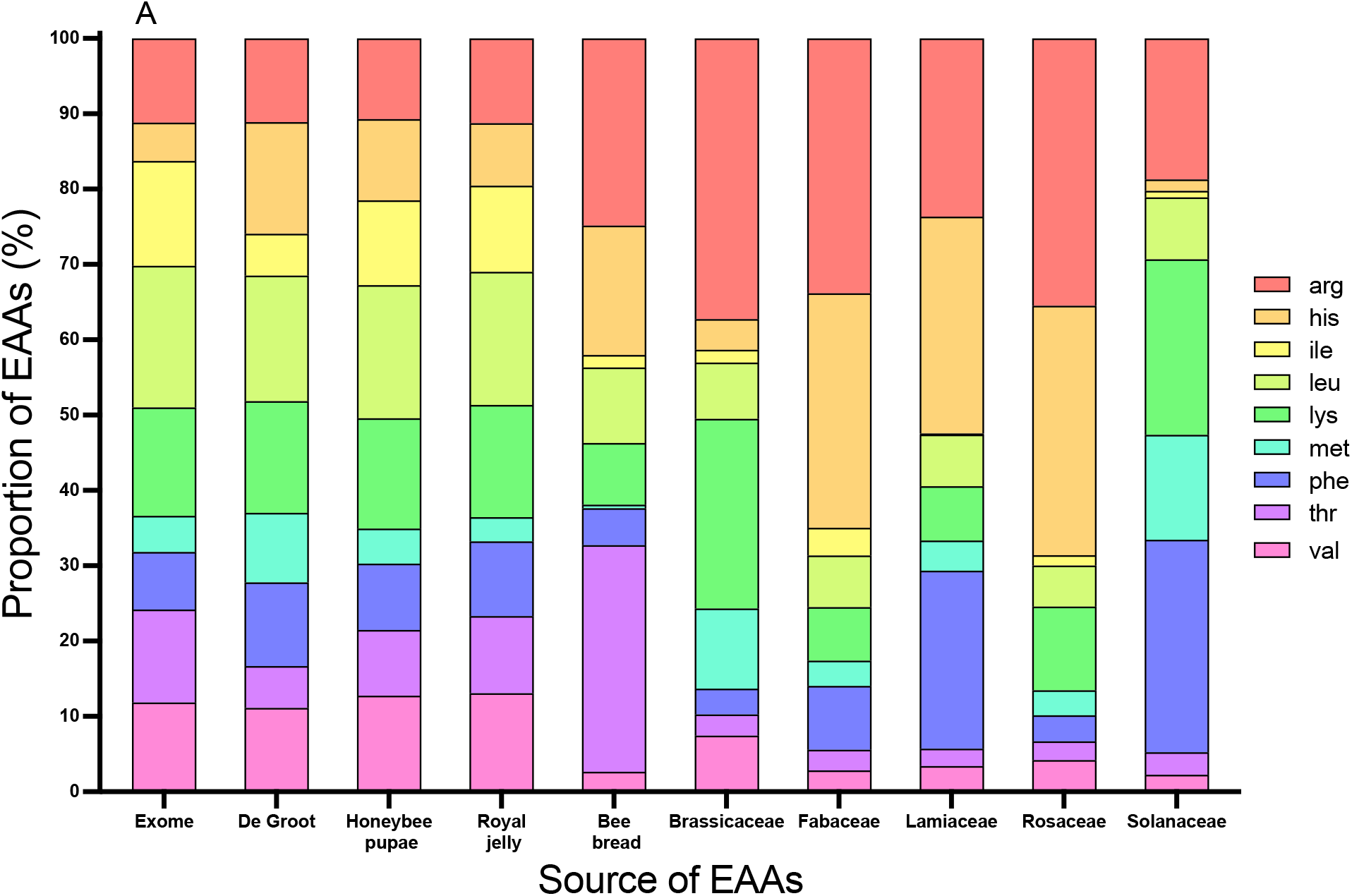
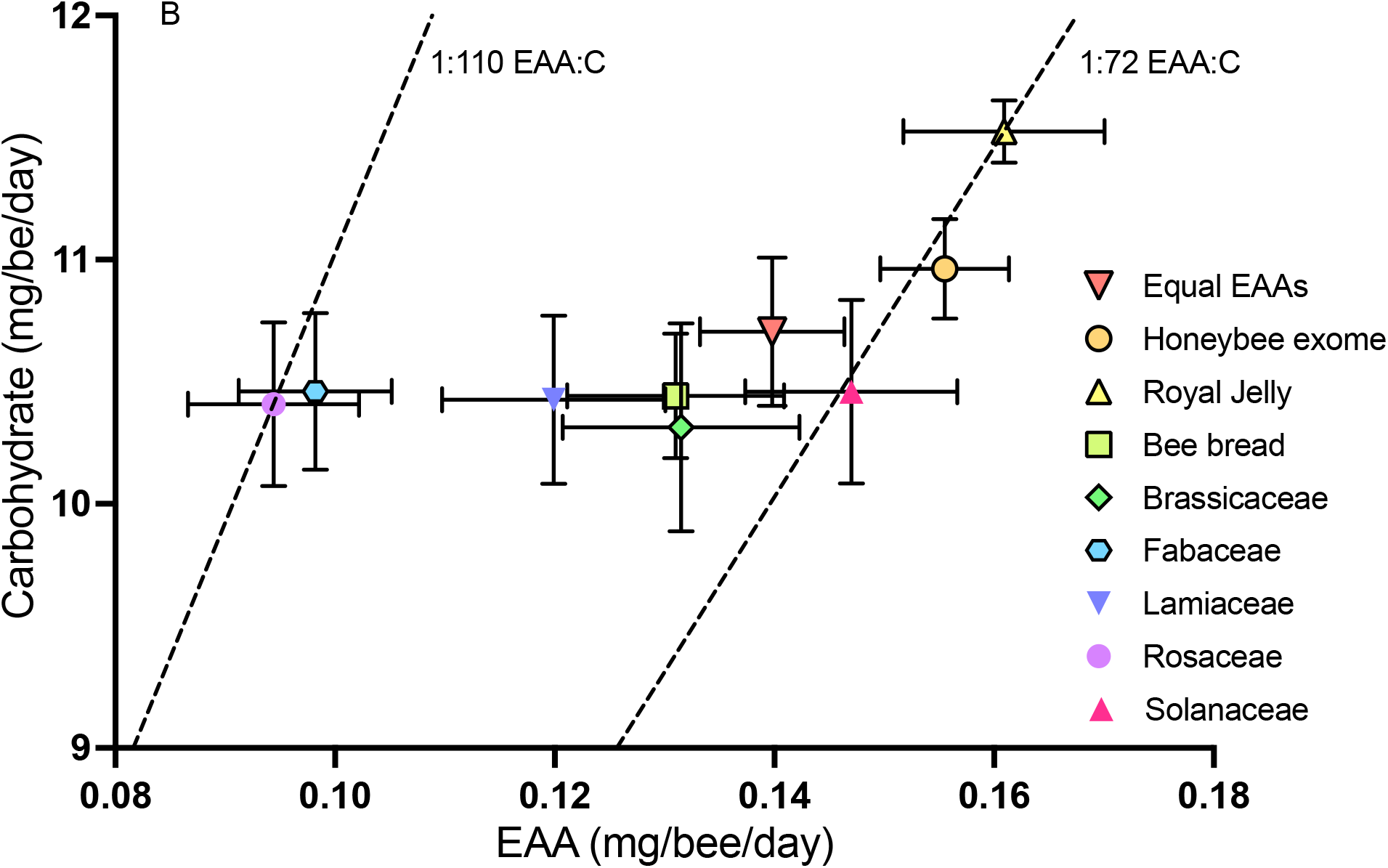
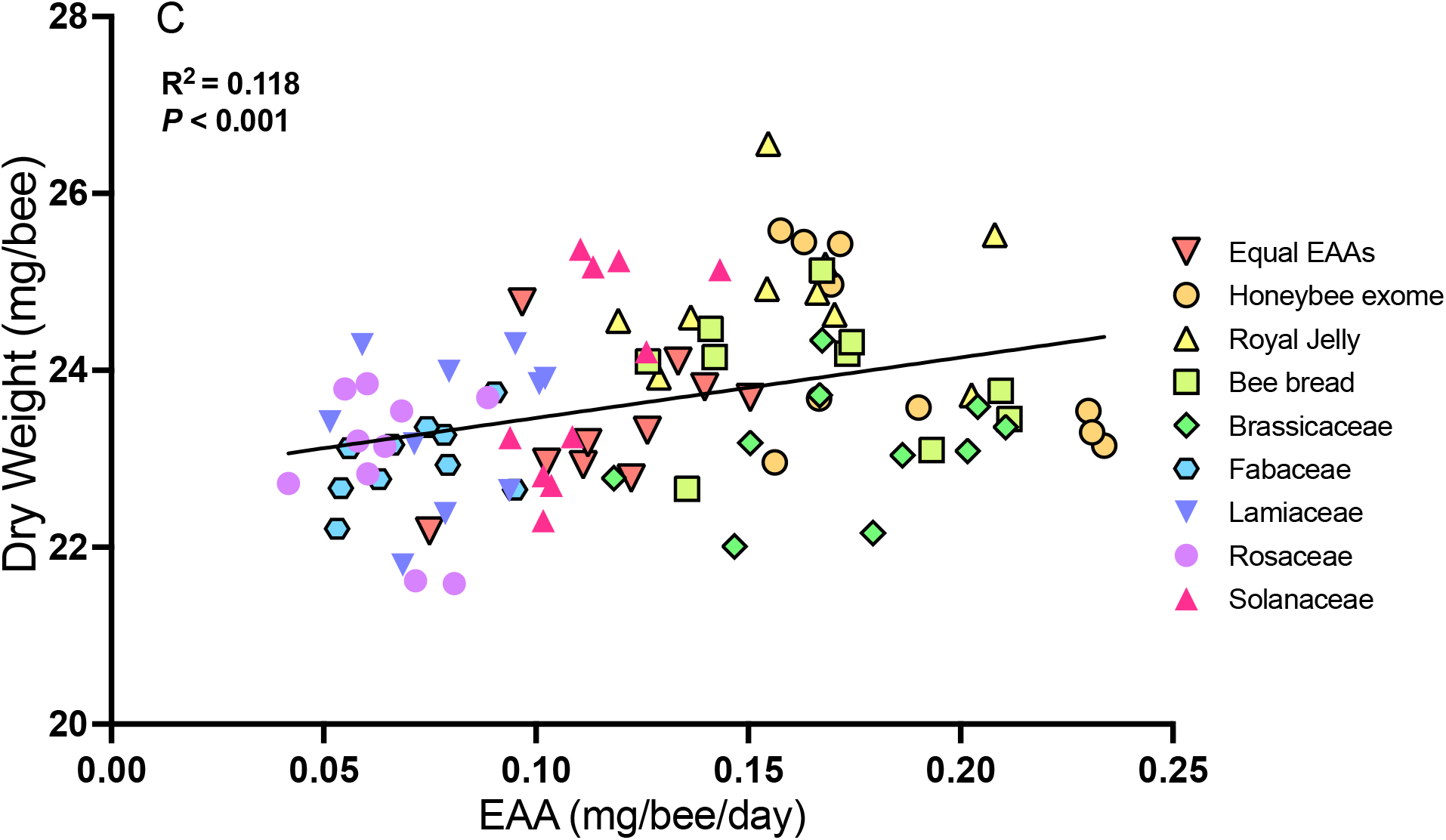
Essential amino acid (EAA) composition, honeybee EAA and carbohydrate consumption, and dry weight of bees after feeding on diets with EAA profiles matching five UK plant families, bee bread, and honeybee tissues. **A**. Proportion of essential amino acids in bee tissues (honeybee exome, De Groot, honeybee pupae, and royal jelly) and those measured in stored pollen (mixed source/bee bread), and pollen of five UK plant families. **B**. Average daily intake of essential amino acids and carbohydrate. EAA intake on Royal Jelly, Equal EAA and Solanaceae diets did not differ from Honeybee Exome (*LSD P’s* ≥ 0.099). In all other treatment diets, bees consumed less EAA than bees fed the EAA profile of honeybee exome (*LSD P’s* ≤ 0.016). **C**. Dry weight of surviving honeybees after 10 days of feeding on EAA-defined diets.

The bees chose to eat proportionally more EAAs when the diets matched honeybee tissues or products (Figure 1B. MANOVA, EAA: F_8,200_ = 7.38 *P* < 0.001). The mean daily amount of carbohydrates consumed was not significantly different among treatments (MANOVA, C: F_8,200_ = 1.11 *P* = 0.357); on average, bees ate ∼10.7 mg of carbohydrate in all treatments. As the quantity of EAAs varied as a function of diet, each treatment group consumed different ratios of EAA:C. Bees fed with the Exome, Royal Jelly, Equal EAA, or Solanaceae treatments achieved a ratio of EAA:C of ∼1:72. However, bees fed the other diet treatments achieved the following ratios: 1:78 (Brassicaceae), 1:80 (Bee Bread), 1:87 (Lamiaceae), 1:107 (Fabaceae), and 1:110 (Rosaceae). Remarkably, the honeybees ate 1.7 times less EAA when fed the Fabaceae and Rosaceae diets (Figure 1B) compared to the Bee Exome diet. (Note: prior to the start of this experiment, we also tested Exome, Pupal hydrolysis and the EAA profile predicted by de Groot (1953) (Table S5). The bees’ feeding behaviour was not significantly different on any of these treatments, so all experiments were conducted with the Exome treatment (Figure S2).

We also measured the rate of mortality during the experiment and weighed a subset of the surviving honeybees. Approximately 90% of the bees that started in the experiment survived the 10-day treatment; the rate of mortality over the period was not significantly affected by diet (Coxreg, 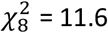, *P* = 0.171). The dry weight of the bees was a function of the EAAs consumed in diet (Figure 1C, linear regression, R2 = 0.118, *P* < 0.001). Bees that had been fed with the EAA profile of Royal Jelly or Exome had a greater total dry weight than bees fed with the other diets (Figure 1C, ANOVA, F_8, 81_ = 5.56, *P* < 0.001). Each section of the bee’s body was affected by diet (Figure S3, MANOVA, Abdomen: F_8,81_ = 4.21, *P* < 0.001; Thorax: F_8,81_ = 5.55, *P* < 0.001; Head: F_8,81_ = 6.31, *P* < 0.001). However, the protein content of bee thoraxes was not significantly affected by the EAA treatment that they were feeding on (Figure S4, ANOVA, F_8,81_ = 0.957, *P* = 0.476).

### Experiment 2: Branched chain amino acids in pollen conflict with arginine and histidine

To verify which of the EAAs were contributing to potential differences in food consumption and performance, we entered the EAA profiles of all the diets into a factor analysis (Table 1, Figure 2A, 2B, 2C). Three factors with eigenvalues > 1 were selected which accounted for 87.8% of the variation in the data (Table 1, Figure 2A, 2B, 2C). The first factor described a strong positive correlation between the three BCAAs. The quantities of these EAAs were inversely related to arginine and histidine. The second factor described a positive correlation between lysine and methionine; these EAAs were inversely related to threonine. The final factor best described the proportion of phenylalanine in each diet.

We designed an experiment in which we constructed diets that represented the proportions of the EAAs described by each factor in experiment 1. For example, Factor 1A (Figure 2A) represented a diet in which the BCAAs were proportionally higher than histidine and arginine; Factor 1B had elevated histidine and arginine, and reduced BCAAs. In each factor, the EAAs that did not have high weightings were held at a constant proportion of the diet (Table S5). In the Factor 1 diets, the bees fed the Factor 1A diet ate 1.24 times more EAAs and 1.07 times more carbohydrate compared to those fed the Factor 1B diet (Figure 2D, Generalised linear model, EAA: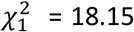, *P* < 0.001, C: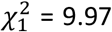, *P* < 0.001). Factor 2A and 2B representing the relationship of lysine and methionine relative to threonine did not significantly affect food intake (Figure 2E, Generalised linear model, EAA: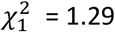, *P* = 0.256, C: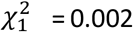, *P* = 0.965). Factor 3, representing phenylalanine relative to all other EAAs also had a significant influence on food consumption (Figure 2F, Generalised linear model, EAA: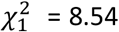, *P* = 0.003), as bees fed with elevated phenylalanine ate 1.23 times more EAAs than those subject to diets with low levels of phenylalanine. However, the proportion of phenylalanine in the diet did not affect their carbohydrate intake (Figure 2F, Generalised linear model C: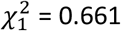, *P* = 0.416). None of the diets we tested here significantly influenced bee survival in the experiment (Factor 1: Coxreg, 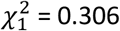, *P* = 0.58; Factor 2: Coxreg, 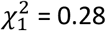, *P* = 0.597; Factor 3: Coxreg,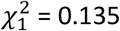, *P* = 0.713).

**Figure 2.**
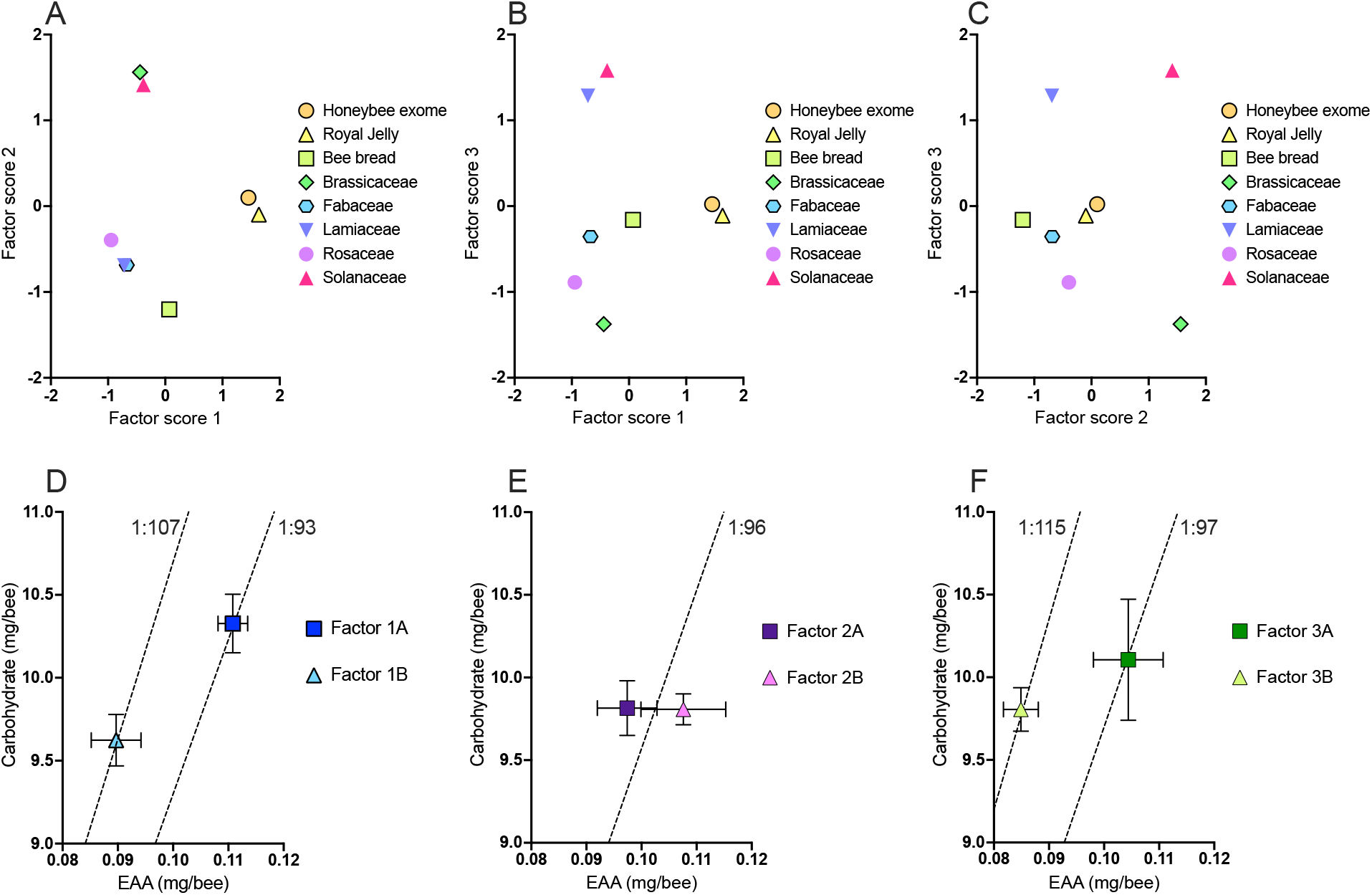
Principal components from a factor analysis of honeybee tissues, bee bread, and the pollen of five UK plant families, and the resulting EAA and C feeding behaviour of honeybees fed diets representing both inverse and positive correlations of EAAs associated with each of three factors. **A**. Factor 1 represented a strong positive correlation between the presence of the three branched chain amino acids (leucine, isoleucine and valine) inversely correlated with the presence of arginine and histidine. Honeybee tissues (honeybee exome and royal jelly) are clearly separated from the pollen profiles representing 5 different plant families. **B**,**C**. A second factor represented a positive correlation between the presence of lysine and methionine with an inverse relationship to the presence of threonine. This separated Brassicaceae and Solanaceae most strongly from bee bread. A third factor represented the proportion of phenylalanine relative to all other EAAs. The EAA profile of Lamiaceae and Solanaceae different most greatly from the profile of Brassicaceae. **D**. Elevated BCAAs (Factor 1A EAAS) drove bees to modify intake of both EAA and C, to a more EAA-biased intake of 1:93 EAA:C. However, when BCAAs were supressed relative to arg and his (Factor 1B EAAs), bees consumed a carbohydrate biased intake of 1:107 EAA:C. **E**. Elevation or reduction of lysine, methionine or threonine relative to the other 7 EAAS did not influence the way in which honeybees consumed EAA and C and in both Factor 2A and 2B treatments bees achieved similar EAA:C intakes of 1:96. **F**. Elevation of phenylalanine in proportion to all other EAAs (Factor 3A) caused bees to increase their total EAA intake relative to diets where phenylalanine was proportionally lower than all other EAAs (Factor 3B) resulting in a more EAA-biased intake of 1:97 EAA:C relative to 1:115 EAA:C.

### Experiment 3: The ratio of BCAAs to histidine is used to regulate the intake of EAAs in diet

Of the BCAAs, leucine is the main signal in animal cells of the presence of protein, as it activates mTORC1 (Lynch et al., 2000; Gould et al., 2023; Reifenberg and Zimmer, 2024). For this reason, we hypothesized that the patterns of dietary intake that we observed could be driven by the quantities of leucine in diet as previously observed (reviewed by Pedroso et al., 2015). To test this, we created three diets where the ratio of leucine varied relative to at least one of the other EAAs we observed to account for the patterns of feeding in Figure 2 above. As predicted, the proportion of histidine in diet strongly influenced EAA feeding (two-way MANOVA, EAA pair × ratio, EAA F_6,108_ = 2.54, *P* = 0.024). When bees were fed diets where the proportion of histidine exceeded that of leucine, they significantly reduced their protein feeding (Figure 3A, *P* ≤ 0.022). In fact, the treatment diet in which histidine was greatest relative to leucine, bees shifted their EAA:C intake to 1:110 compared to 1:70 in the treatment in which leucine was highest relative to histidine. However, the proportion of leucine relative to arginine (Figure 3B) or lysine (Figure 3C) did not influence honeybee EAA or carbohydrate feeding, and in both experiments, bees converged on similar intakes of 1:83 and 1:82 EAA:C on leucine-to-arginine and leucine-to-lysine experiments, respectively (*P*’s > 0.05). Our experiments clearly indicated that the pattern we observed in Figure 2A, could be driven by the relationship between the relative amounts of leucine-to-histidine that was arising in part because of histidine’s role in suppression of feeding. To identify if this was specific to the leucine-to-histidine ratio, we constructed a second experiment with diets where the ratio of histidine varied relative to the other two BCAAs (isoleucine and valine) and lysine. As before, we found that the quantity of EAA consumed in diet was a function of the ratio of histidine to any of the BCAAs (Figure 3D, Figure 3E, two-way MANOVA, EEA pair × ratio, EAA, F_6,108_ = 5.37, *P* <0.001). Interestingly, the proportion of histidine to the BCAAs also caused bees to modify their carbohydrate intake (two-way MANOVA, EEA pair × ratio, C, F_6,108_ = 2.83, *P* = 0.014). Like the trend observed in histidine-to-leucine experiments, bees shifted their EAA to C intake from 1:70 EAA:C to 1:118 EAA:C (Figure 3D) and 1:65 EAA:C to 1:107 EAA:C (Figure 3E) in the histidine-to-valine and histidine-to-isoleucine experiments, respectively. Histidine did not influence the amount of EAAs consumed for diets where histidine was varied relative to lysine (Figure 3F, *P’s* > 0.05); bees fed with the high lysine-low histidine diet, however, consumed more EAA than any of the other diets (*P’s* < 0.001). To understand if taste of histidine was driving nutrient regulation, a subset of bees was used in an experiment to test their likelihood of drinking a solution that matched the equal EAA control diet, the low leucine-high histidine, and high leucine-low histidine diets. Bees were equally likely to drink from all three test solutions used (Figure S5, ANOVA, F_2,39_ = 0.50, *P* = 0.610).

**Figure 3.**
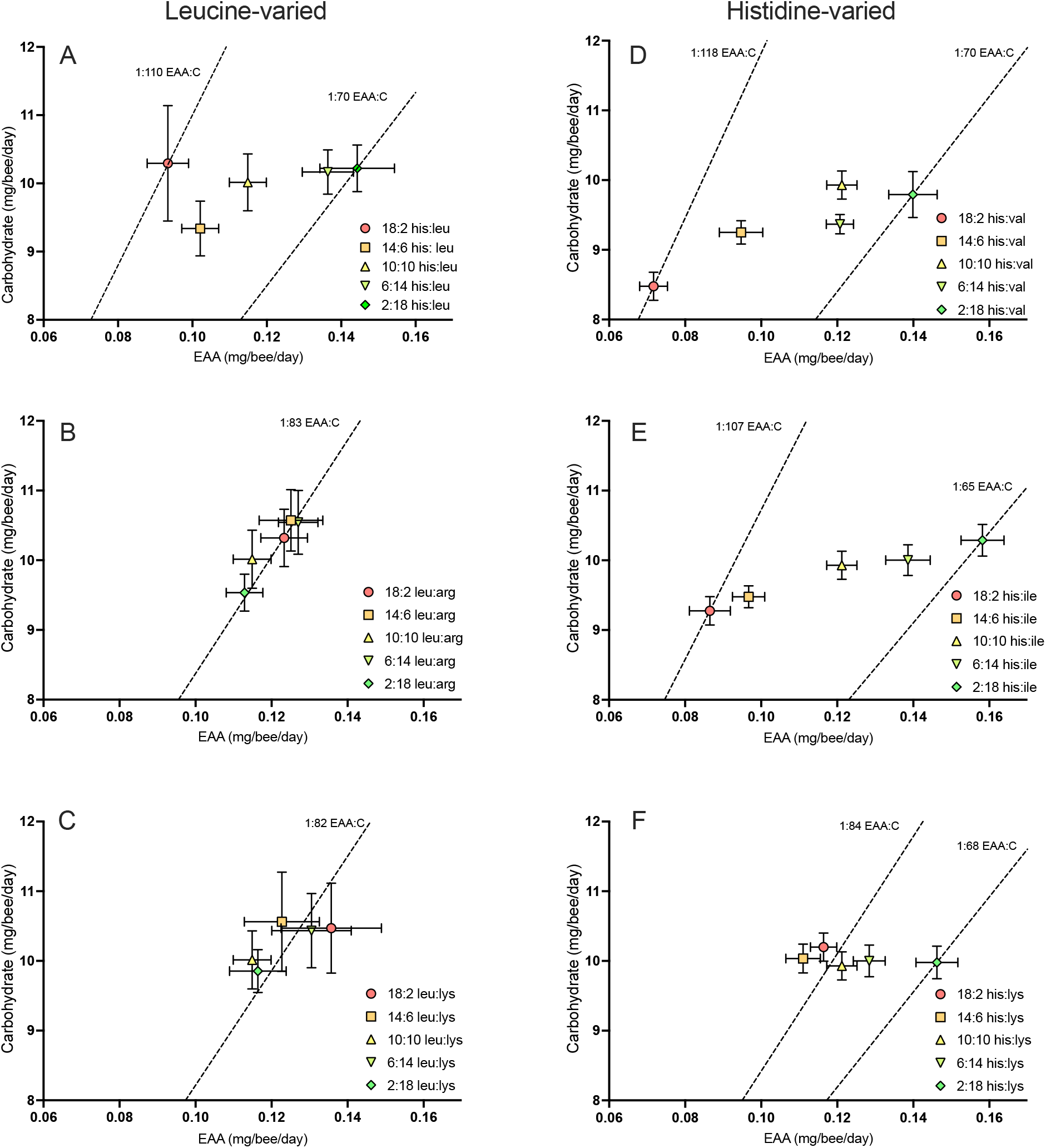
Essential amino acid and carbohydrate balancing is constrained by the proportion of the BCAAs relative to histidine in diet. **A**. Manipulating the proportion of leucine relative to histidine from 2:18 to 18:2 induces a 1.6-fold increase in EAA consumption relative to carbohydrate. **B**. Manipulating the proportion of leucine relative to arginine in diet does not influence the way in which honeybees balance their EAA:C intake and in all treatments bees converge on an intake of 1:83 EAA:C. **C**. Manipulating leucine relative to lysine does not influence honeybee EAA:C balance and bees converge on an intake target of 1:82 EAA:C. **D**. Manipulating the proportion of histidine relative to valine affects honeybee EAA:C balancing. In situations where histidine outweighs valine, bees undereat EAA achieving an intake target of 1:118. When valine exceeds the proportion of histidine in diet, bees eat more EAAs and achieve and intake target of 1:70 EAA:C. **E**. The proportion of histidine relative to isoleucine in diet affects EAA:C balancing. When histidine is present at 18:2 relative to isoleucine, bees undereat EAAs and achieve an EAA:C intake of 1:107. However, when isoleucine exceeds the proportion of histidine in diet (2:18 his:val), bees consume more EAAs and achieve an EAA:C intake of 1:65. **F**. High presence of histidine in diet relative to lysine does not cause bees to undereat EAAS. However, the presence of high lysine relative to histidine causes bees to increase their EAA:C intake from 1:84 to 1:68 EAA:C.

## Methods

### Animals

Frames of sealed brood were taken from honeybee (*Apis mellifera*) colonies at the apiary, Oxford University, Wytham, UK. Brood frames were housed in a ventilated box and kept in a climate-controlled incubator at 34 °C and 60% relative humidity. Within 24 h of emerging, female bees from all frames were brushed into a ventilated box and allowed to form a homogenous population for 5 min. Cohorts of 30 bees were then collected and housed in Perspex feeding cages with six holes for feeding tubes (Paoli et al., 2014; Stabler et al., 2021). Bees were provided with solid diets with *ad libitum* access to water via modified 2 ml microcentrifuge tubes with 4 × 2 mm holes drilled along their length. Solid diets were also provided in 2 ml microcentrifuge tubes, but they were modified so that they had 3 × 5 mm holes drilled along one side. Feeding cages with bees were kept in climate-controlled chambers (HPP570, Memmert GmBH + Co. KG.) at 34 °C and 60% relative humidity for either 5 or 10 days, depending on the experiment. Mortality was measured daily, and any dead bees were removed from the feeding cages.

### Diets

Solid diets were formulated for all experiments using powdered sugar, 80% invert syrup, a lipid source, maltodextrin and a blend of reagent grade EAAs (Table S4). The EAA:C composition of all diets was 1:25 EAA:C w/w. This ratio was chosen because nurse-aged honeybees can balance their intake of both EAA and C macronutrients on diets in this range (Paoli et al., 2014). Carbohydrate was maintained at 75% of the end diet and free amino acids were incorporated at 3% in treatment diets. Control diets did not contain any EAAs and this nutrient space was replaced with maltodextrin to ensure the proportions of all diets equated to 100% (Table S4). Diets contained 2% fat providing an α-linolenic : linoleic acid ratio of 1:1 using a combination of flax oil and soy lecithin and water (Sereia et al., 2013; Vaudo et al., 2014; Arien et al., 2015; 2018; Stabler et al., 2021).

### Pollen HPLC analysis

Pollen EAAs were quantified from UK flowering plant families using the methods described in Stabler et al., 2018. Pollen was subjected to an HCL microwave-assisted hydrolysis at a proportion of pollen to 6M HCL of 1:100 (mg:µl). Pollen was microwaved for 20 min at 900W in a domestic microwave. After microwaving, samples were allowed to cool before removing from the microwave. Tubes were then opened and placed in a heat block at 100°C until dry. Samples were resuspended in UHPLC gradient grade water and vortexed for 15 min before centrifuging at 13,249 *g* for 30 min. Supernatant was then removed with a sterile 1 ml syringe and then filtered through a 0.45 µm syringe tip filter. The filtered sample was then centrifuged for a further 10 min before analysis with UHPLC (Stabler et al., 2018). The amino acid profile of bee bread was estimated based on the findings reported in Donkersley et al., (2017). Fifty honeybee colonies located in the North-west of England, UK were sampled for stored pollen (bee bread). Their amino acid composition was quantified as in Stabler et al., (2015; 2018) and their average EAA composition was used to represent the EAA values of bee bread in our feeding experiments (Donkersely et al., 2017).

### Experiment 1

The first experiment was designed to test how the measured amino acids from pollen, bee bread, bee tissues, and bee products (Table S2) affected EAA and carbohydrate balancing of newly emerged, adult worker honeybees. The honeybee exome values were derived by calculating the proportions of EAAs from the total number of counts of each EAA in the honeybee exome extracted from the honeybee genome (Bee Base). Diets were made as described above, where the 3% of amino acids were comprised of the relevant treatment groups’ amino acid profiles (Table S3), paired with an amino acid-free control diet. Bees had *ad lib* access to one treatment and a control diet for 10 days. Diets were weighed and replaced every 24 hr. Each treatment was replicated with 20 cohorts of 30 newly emerged bees. After 10 days of feeding, animals were euthanised by freezing and 10 remaining animals from 10 cohorts were dissected into head, thorax, and abdomen before drying to constant mass, at which point, dry weight was recorded. Total nitrogen was measured from the pooled sample of 10 dissected thoraxes at Elemental Microanalysis (elementalmicroanalysis.com) using the method described in Stabler et al., (2018). Total nitrogen was used to estimate protein content by using the standard multiplication factor of 6.25.

### Experiment 2

After observing that bees modified their EAA:C balancing due to the EAA composition of their diet, we designed a series of experiments to explore how the relative elevation or reduction of certain EAAs affected nutrient balancing. We performed a principal components analysis (PCA) on the EAA composition of the treatment diets in experiment 1 (Table 1). Experiments were designed such that the amino acids represented by each of the factors produced by the PCA were elevated or reduced relative to how they were related in the loading on the factor. Manipulations to amino acid composition were made in 20% of the amino acid nutrient space. All other EAAs not associated with the factor were incorporated at the 80% of the amino acid nutrient space (Table S6). This resulted in three diet pairs; Factor 1A represented elevated isoleucine, leucine, and valine, relative to arginine and histidine, whereas 1B represented reduced isoleucine, leucine, and valine, relative to elevated arginine and histidine. Factor 2A represented elevated lysine and methionine relative to reduced threonine and 2B represented reduced lysine and methionine and elevated threonine. Factor 3A represented elevated phenylalanine relative to the other nine EAAs, whereas 3B represented reduced phenylalanine relative to the other nine EAAs (Table S6).

**Table 1.** Principal components analysis of pollen from five UK families of plants.

|  | Factor |  |  |
| --- | --- | --- | --- |
|  | 1 | 2 | 3 |
| Eigenvalue | 3.79 | 2.85 | 1.27 |
| % variance | 42.1 | 31.6 | 14.1 |
| Amino acid |  |  |  |
| arg | **-0.822** | -0.003 | 0.556 |
| his | **-0.655** | -0.633 | -0.111 |
| ile | **0.901** | -0.275 | 0.098 |
| leu | **0.967** | -0.22 | -0.116 |
| lys | 0.287 | **0.865** | 0.364 |
| met | 0.087 | **0.985** | 0.062 |
| phe | -0.056 | 0.548 | **-0.824** |
| thr | 0.261 | **-0.534** | -0.095 |
| val | **0.88** | -0.136 | 0.314 |

### Experiment 3

A final series of experiments were designed to carefully test the effect of elevating and reducing specific EAAS relative to one other EAA. In these experiments, only two EAAs were manipulated, meaning that all other EAAs could be maintained at a constant rate of inclusion of 10% of the amino acid nutrient space (Table S7). Six amino acid combinations were used in which the two target EAAs were incorporated at either 18:2, 14:6, 6:14 or 2:18 based on their percentage contribution to the amino acid nutrient space. A control treatment of 10:10 was also used for each comparison (Table S7). In order to determine if the taste of the amino acids was driving any recorded changes in EAA to C balancing, a subset of adult female honeybees was used in a taste assay. The amino acid composition of equal EAAs, 18:2 leu:his and 2:18 leu:his were used as tastants. Amino acids were dissolved into solution at 1.37% and sucrose was dissolved at 34.23%, so that amino acids and carbohydrate were at the ratio of EAA:C that was used in solid diets (1:25). Adult female bees were restrained in modified 1 ml pipette tips and fed to satiety on 1M sucrose before being left for 18 hours. Bees were then divided and assigned to one of the treatment groups. The antennae of bees were stimulated with a droplet of 1M sucrose to elicit a proboscis extension. Once the proboscis was extended, bees were presented with a droplet of the assigned treatment solution. The feeding response of the bees was recorded as positive or negative based on whether the bee could be observed to actively be drinking by movement of the proboscis, or whether the droplet was consumed.

### Statistical analysis

EAA composition of bee tissues compared to bee bread and hand-collected pollen was compared using a canonical discriminant analysis (CDA). The amino acid profiles of treatments used in experiment 1 were applied to a factor reduction using principal components analysis (PCA). Nutrient consumption was calculated using the daily change in weight of diet and dividing the value by the number of live bees in the box over the same 24 h period. Changes in weight of evaporation controls were used to adjust consumption values. In experiment 1, average daily nutrient intake of EAA and C were compared using multivariate analysis of variance (MANOVA) with *LSD post hocs* used to compare other treatment intakes against the bee tissue treatment. Dry weight of bee head, thorax and abdomen were compared using a one-way ANOVA and pairwise comparisons were made using Šidák’s *post hoc*. Nutrient intake of each treatment in experiment 2 were compared using generalised linear models. Diet consumption of experiment 3 were compared using a two-way ANOVA with treatment and diet as predictors. Nutrient intake was compared using MANOVA with Sidak *post hocs* for pairwise interactions. A cox regression was applied to survival for all three experiments. Thorax protein content and drink response in the taste assay were compared with one-way ANOVA. All statistical analyses were carried out using SPSS V18.9.

## Discussion

Many studies have investigated how protein impacts food intake in animals, but none have shown that its influence is a result of the relative quantities of EAAs. Our work confirms that the regulation of protein by animals depends on the BCAAs in food, but it also shows that the impact of these compounds on feeding depends on the quantities of other EAAs such as histidine. Our novel study shows that pollen protein can be suboptimal for bees in a way that depends on its EAA profile.

An important discovery in our data was the fact that pollen protein with the greatest mismatch in EAAs to the profile found in honeybee tissue caused bees to eat less food, gain less body mass and consume proportionally more carbohydrates (e.g., 1:110 EAA:C vs 1:72 EAA:C). Specifically, we discovered that when the proportion of BCAA-to-histidine in diet was low, bees ate less EAA and proportionally more carbohydrates. Furthermore, specific deficiencies of individual amino acids (i.e. phenylalanine) also shifted macronutrient balancing away from EAAs towards carbohydrates. Importantly, our data show that the relationship between BCAAs and histidine in diet is a key factor responsible for ‘protein leveraging’: the condition whereby animals overconsume carbohydrates relative to protein at a cost to their fitness and survival (Simpson and Raubenheimer, 2005). Our experiments also imply that bees cannot simply eat more pollen to reach nutritional homeostasis for all their EAAs if they are restricted to eating pollen from the Rosaceae or Fabaceae because of the extreme imbalance in histidine relative to the BCAAs in these families’ pollen protein.

In Drosophila, larvae fed with the EAA profile represented in the Drosophila ‘exome’ eat more, develop faster and are more apt to survive (Piper et al., 2017). EAAs in diet also matter for reproduction: adult female Drosophila or copepods produce fewer eggs when fed diets where the EAA profile deviates from their exome or their tissues (Guisande et al., 1999, 2000; Ma et al., 2022). Mouse studies have also recently revealed that diets with amino acids closely resembling the mouse exome, at static total protein concentrations, afford mice increased growth rate and improved feed conversion efficiency relative to diets mis-matched in their amino acids, relative to their exome, with improved lean body mass and higher sperm production of male mice (Wu et al., 2024). Our study observed increased mass of nurse-aged female workers but as worker bees are not reproductive, we have no data to corroborate these other metrics. Future studies should explore the role of dietary amino acids on queen honeybee performance. Restricting individual amino acids such as isoleucine also improves *Drosophila* resistance to nicotine poisoning (Fulton et al., 2022; Fulton et al., 2024) far surpassing the effect of manipulating total protein. This strongly suggests that protein quality is an important factor when considering an animal’s resistance to environmental stressors. Here, we have shown that nurse-aged worker honeybees modify their protein and carbohydrate feeding behaviour in response to the essential amino acid composition of diet. It is likely that variation in both total protein and quality of protein consumed affect honeybees’ resistance to environmental stressors and that this is limited by the plant species pollen that they can forage on. It is likely that manipulating individual amino acids, particularly the branched-chain amino acids, modifies protein-signalling pathways such as mTORC1 and GCN2 (Fulton et al., 2022). Our findings clearly demonstrate that the presence of histidine in diet relative to the three branched-chain amino acids influences honeybee protein-feeding. It is likely that this will also impact protein-signalling pathways in honeybees, but this was not explored in our study. Indeed, honeybees fed high levels of dietary amino acids have reduced lifespans and low expression of Sir2, compared to bees with increased lifespan when fed low levels of dietary essential amino acids (Paoli et al., 2014). In our study, we compared the honeybee ‘exome’ to the predictions for EAA of de Groot (1953) and found that they are quite similar and that feeding behaviour on diets composed of these predicted EAA ratios were not significantly different. Interestingly, the EAAs rendered by the digestion of royal jelly are also a close match to what is needed by bee tissues. We predict that honeybees have evolved to create glandular secretions which are the perfect food for their larvae, providing them with the exact ratios of EAAs that maximize growth. Other bee species, however, feed pollen directly to larvae and cannot mitigate nutritional deficiencies in the pollen except to provision a mix of available pollens.

It is well-known that generalist bees forage for pollen from several plant sources to create a mixture, for nest provisioning in solitary species, and ‘bee bread’ in eusocial species (Donkersley et al., 2017; Bednarska et al., 2022). Very few solitary bee species rely on the pollen of one or a few plant species to provision their larvae (Danforth et al., 2019). Our data clearly show that honeybees create a ‘bread’ by mixing pollen sources which is at least as good as the best mono-familial plant source (e.g., Solanaceae). This strategy for pollen foraging is likely to be beneficial not only because it provides missing nutrients, but it also permits bees to mix protein sources in a way that allows them to avoid constraints caused by overeating histidine relative to any of the BCAAs.

Most plant food sources are likely to have EAA profiles that deviate from what an animal needs in predictable ways. An important finding from our research is the fact that the pollen of some plant species (e.g., Roseacea or Fabaceae) is nutritionally detrimental to bees if consumed in the absence of other proteins because of their excess arginine and histidine. Other studies have also observed that pollen sometimes has excess histidine and arginine (Hocherl et al., 2012; Jeannerod et al., 2022). The presence of arginine and histidine were positively correlated in our data set and inversely correlated with the BCAAs (Figure 1A, Table 1). Other studies of the EAA profiles of plant proteins have also observed the inverse relationship between arginine and BCAAs such as leucine (MacArthur et al., 2021, Hertzler et al., 2020). The challenge of dealing with excess arginine (and potentially histidine) to acquire sufficiency in the BCAAs, therefore, is likely to be a problem for all animals that rely on plant proteins.

Histidine is one of the amino acids that is required in the smallest quantities in animals (Brosnan and Brosnan, 2020), and based on the exome we calculated, it is also one of the EAAs required in the lowest quantity by honeybees (and drosophila, Piper et al., 2017) (Figure 1). Furthermore, bees and fruit flies avoid histidine as a free amino acid in food (Hendriskma and Shafir, 2014; Aryal and Lee, 2021). In our work, we did not see a rejection of food as a function of the amount of histidine in the diet (Figure S7), but histidine was mixed with all the other EAAs and so might not have been detectable by taste (Figure S7). For this reason, we expect that the suppression of feeding in our experiments occurred because of post-ingestive feedback caused by the stress of over consumption of histidine.

Previous research has shown that rats fed with high levels of histidine in diet relative to other EAAs eat less food and also gain less body fat (Sheiner et al., 1985; Kasaoka et al., 2004; Endo et al., 2010). The mechanism of suppression in mammals occurs as a result of the conversion of histidine into histamine in the hypothalamus; elevated histamine activates histaminergic receptors (H1) in the brain that are involved in the control of food intake (for a review see Ishizuka and Yamatodani, 2012). One study in rats showed that histidine injection suppressed total feeding by as much as 60%, but it did not impact the relative amount of protein or carbohydrate the rats ate (Sheiner et al., 1985). Histamine is also perceived as an aversive tastant stimulus in insects (Aryal and Lee, 2021).

In mammals, overconsumption of histidine results in increased histamine production. Similarly, in insects increased histamine production causes stress on the Malpighian tubules and elevated histamine excretion (Gaire et al., 2022). In fact, blood or sap sucking insects like tsetse flies, bedbugs, and aphids, actively excrete large amounts (up to 35% wet wt) of arginine and histidine (Bursell, 1965; Prosser and Douglas, 1991; Moriyama and Fukatsu, 2022). Aphids often feed on sap with a free amino acid profile that is highly mismatched to the EAA profile of their tissues (Moriyama and Fukatsu, 2022). To compensate for inadequate supplies of the BCAAs, these insects, as well as the pacific beetle cockroach, have adopted microbial symbionts which provide these particular EAAs (Moriyama and Fukatsu, 2022, Ayayee et al., 2024), but they also excrete arginine and histidine when they are present in food in excess (up to 44% of total excreted amino acids and 72% of amino-nitrogen content, Prosser and Douglas, 1991). The excretion of histidine and arginine are likely to be a compensatory mechanism in these insects that permits them to overcome the costs of ingesting excess histidine relative to BCAAs. Bees may also be excreting these compounds when fed diets like the one we created based on Rosaceous or Fabeaceous pollen, but we did not test this.

Changes in land use have resulted in a loss of diversity of pollen sources for bees (Gillespie et al., 2022; Peters et al., 2022; Birkenbach et al., 2024). Our data imply that the reduction in flowering plant species as pollen sources could potentially increase the potential for EAA malnutrition in bees and other pollinators. Based on studies in Drosophila, we expect that the mismatches that we observed for adult bees would also impact larval growth and development (Piper et al., 2017). For this reason, our data indicate that land managers should consider the EAA profile of floral pollen for habitat amelioration schemes for protecting bee biodiversity, to provide flowering plant species with the best match to the EAAs in pollinator tissues.

## Conflict of interest statement

The authors Daniel Stabler, Sharoni Shafir and Geraldine A. Wright are shareholders in Apix Biosciences, a bee-nutrition university spin-off registered in Belgium. The authors declare that they have no other competing interests.

### Acknowledgements

The authors would like to thank Mathilde Baude, Jane Memmott, Adam Kent, and Rob Wood for their help in pollen collection and analysis. We thank Rui S.F. Gonçalves for beekeeping. This research was funded in whole or in part by the BBSRC grant numbers *BB/P007449/1, BB/T014210/1 to* GAW and SS. For the purpose of Open Access, the author has applied a CC BY public copyright licence to any Author Accepted Manuscript (AAM) version arising from this submission.

## Author contributions

The authors contributed to the manuscript as follows (Daniel Stabler (DS), Jennifer A. Chennells (JAC), Eileen F. Power (EFP), Jennifer Scott (JS), Giorgia Perri (GP), Phillip K. Donkersely (PKD), Sharoni Shafir (SS), and Geraldine A. Wright (GAW)). Conceptualisation: DS, JAC, SS, GAW. Methodology: DS, JAC, GAW. Data collection: DS, JAC, EFP, JS, GP, PKD. Data analysis: DS, GAW. Writing: DS, GAW. Editing: DS, GAW, EFP, GP, SS. funding acquisition: GAW, SS.

## References

Arien, Y., Dag, A. and Shafir, S. (2018). Omega-6: 3 ratio more than absolute lipid level in diet affects associative learning in honey bees. Frontiers in Psychology 9, 1001.

Arien, Y., Dag, A., Zarchin, S., Masci, T. and Shafir, S. (2015). Omega-3 deficiency impairs honey bee learning. Proceedings of the National Academy of Sciences 112, 15761–15766.

Aryal, B. and Lee, Y. (2021). Histamine gustatory aversion in Drosophila melanogaster. Insect Biochemistry and Molecular Biology 134, 103586.

Ayayee, P. A., Petersen, N., Riusch, J., Rauter, C. and Larsen, T. (2024). Enhanced gut microbiome supplementation of essential amino acids in Diploptera punctata fed low-protein plant-based diet. Frontiers in insect science 4, 1396984.

Balnave, D. and Barke, J. (2002). Re-evaluation of the classical dietary arginine: lysine interaction for modern poultry diets: a review. World’s Poultry Science Journal 58, 275–289.

Baude, M., Kunin, W. E., Conyers, S., Davies, N., Gillespie, M. A., Morton, R. D. and Smart, S. M. (2016). Historical nectar assessment reveals the fall and rise of floral resources in Britain. Nature 530, 85–88.

Bednarska, A. J., Mikołajczyk, Ł., Ziółkowska, E., Kocjan, K., Wnęk, A., Mokkapati, J. S., Teper, D., Kaczyński, P., Łozowicka, B. and Śliwińska, R. (2022). Effects of agricultural landscape structure, insecticide residues, and pollen diversity on the life-history traits of the red mason bee Osmia bicornis. Science of The Total Environment 809, 151142.

Birkenbach, M., Straub, F., Kiesel, A., Ayasse, M., Wilfert, L. and Kuppler, J. (2024). Land-use affects pollinator-specific resource availability and pollinator foraging behaviour. Ecology and Evolution 14, e11061.

Brosnan, M. E. and Brosnan, J. T. (2020). Histidine metabolism and function. The Journal of Nutrition 150, 2570S–2575S.

Bursell, E. (1965). Nitrogenous waste products of the tsetse fly, Glossina morsitans. Journal of Insect Physiology 11, 993–1001.

Cook, S. M., Awmack, C. S., Murray, D. A. and Williams, I. H. (2003). Are honey bees’ foraging preferences affected by pollen amino acid composition? Ecological Entomology 28, 622–627.

Dai, Z., Zheng, W. and Locasale, J. W. (2022). Amino acid variability, tradeoffs and optimality in human diet. Nature communications 13, 6683.

Danforth, B. N., Minckley, R. L. and Neff, J. L. (2019). The solitary bees: biology, evolution, conservation. Princeton University Press.

de Groot, A. P. (1953). Protein and amino acid requirements of the honeybee (Apis mellifica L.). Physiol. Comparate et Oecologia 3, 197–285.

Donkersley, P., Rhodes, G., Pickup, R. W., Jones, K. C., Power, E. F., Wright, G. A. and Wilson, K. (2017). Nutritional composition of honey bee food stores vary with floral composition. Oecologia 185, 749–761.

Dufour, C., Fournier, V. and Giovenazzo, P. (2020). Diversity and nutritional value of pollen harvested by honey bee (Hymenoptera: Apidae) colonies during lowbush blueberry and cranberry (Ericaceae) pollination. The Canadian Entomologist 152, 622–645.

Endo, M., Kasaoka, S., Takizawa, M., Goto, K., Nakajima, S., Moon, S.-K., Kim, I.-S., Jeong, B.-Y. and Nakamura, S. (2010). Suppressed fat accumulation in rats fed a histidine-enriched diet. Preventive Nutrition and Food Science 15, 1–6.

Fulton, T. L., Mirth, C. K. and Piper, M. D. (2022). Restricting a single amino acid cross-protects Drosophila melanogaster from nicotine poisoning through mTORC1 and GCN2 signalling. Open Biology 12, 220319.

Fulton, T. L., Johnstone, J. N., Tan, J. J., Balagopal, K., Dedman, A., Chan, A. Y., Johnson, T. K., Mirth, C. K. and Piper, M. D. (2024). Transiently restricting individual amino acids protects Drosophila melanogaster against multiple stressors. Open Biology 14, 240093.

Gaire, S., Principato, S., Schal, C. and DeVries, Z. C. (2022). Histamine excretion by the common bed bug (Hemiptera: Cimicidae). Journal of Medical Entomology 59, 1898–1904.

Ghosh, S., Jeon, H. and Jung, C. (2020). Foraging behaviour and preference of pollen sources by honey bee (Apis mellifera) relative to protein contents. Journal of Ecology and Environment 44, 4.

Gillespie, M. A., Baude, M., Biesmeijer, J., Boatman, N., Budge, G. E., Crowe, A., Davies, N., Evans, R., Memmott, J. and Morton, R. D. (2022). Landscape-scale drivers of pollinator communities may depend on land-use configuration. Philosophical Transactions of the Royal Society B 377, 20210172.

Goul, C., Peruzzo, R. and Zoncu, R. (2023). The molecular basis of nutrient sensing and signalling by mTORC1 in metabolism regulation and disease. Nature Reviews Molecular Cell Biology 24, 857–875.

Guisande, C., Maneiro, I. and Riveiro, I. (1999). Homeostasis in the essential amino acid composition of the marine copepod Euterpina acutifrons. Limnology and Oceanography 44, 691–696.

Guisande, C., Riveiro, I. and Maneiro, I. (2000). Comparisons among the amino acid composition of females, eggs and food to determine the relative importance of food quantity and food quality to copepod reproduction. Marine Ecology Progress Series 202, 135–142.

Hanley, M. E., Franco, M., Pichon, S., Darvill, B. and Goulson, D. (2008). Breeding system, pollinator choice and variation in pollen quality in British herbaceous plants. Functional Ecology 592–598.

Hendriksma, H. P., Oxman, K. L. and Shafir, S. (2014). Amino acid and carbohydrate tradeoffs by honey bee nectar foragers and their implications for plant–pollinator interactions. Journal of insect physiology 69, 56–64.

Hertzler, S. R., Lieblein-Boff, J. C., Weiler, M. and Allgeier, C. (2020). Plant proteins: assessing their nutritional quality and effects on health and physical function. Nutrients 12, 3704.

Hsu, P.-S., Wu, T.-H., Huang, M.-Y., Wang, D.-Y. and Wu, M.-C. (2021). Nutritive value of 11 bee pollen samples from major floral sources in Taiwan. Foods 10, 2229.

Ishizuka, T. and Yamatodani, A. (2012). Integrative role of the histaminergic system in feeding and taste perception. Frontiers in systems neuroscience 6, 44.

Jeannerod, L., Carlier, A., Schatz, B., Daise, C., Richel, A., Agnan, Y., Baude, M. and Jacquemart, A.-L. (2022). Some bee-pollinated plants provide nutritionally incomplete pollen amino acid resources to their pollinators. PLoS One 17, e0269992.

Kasaoka, S., Tsuboyama-Kasaoka, N., Kawahara, Y., Inoue, S., Tsuji, M., Ezaki, O., Kato, H., Tsuchiya, T., Okuda, H. and Nakajima, S. (2004). Histidine supplementation suppresses food intake and fat accumulation in rats. Nutrition 20, 991–996.

Leinonen, I., Iannetta, P. P., Rees, R. M., Russell, W., Watson, C. and Barnes, A. P. (2019). Lysine supply is a critical factor in achieving sustainable global protein economy. Frontiers in Sustainable Food Systems 3, 27.

Lynch, C. J., Fox, H. L., Vary, T. C., Jefferson, L. S. and Kimball, S. R. (2000). Regulation of amino acid– sensitive TOR signaling by leucine analogues in adipocytes. Journal of cellular biochemistry 77, 234–251.

Ma, C., Mirth, C. K., Hall, M. D. and Piper, M. D. (2022). Amino acid quality modifies the quantitative availability of protein for reproduction in Drosophila melanogaster. Journal of insect physiology 139, 104050.

MacArthur, M. R., Mitchell, S. J., Treviño-Villarreal, J. H., Grondin, Y., Reynolds, J. S., Kip, P., Jung, J., Trocha, K. M., Ozaki, C. K. and Mitchell, J. R. (2021). Total protein, not amino acid composition, differs in plant-based versus omnivorous dietary patterns and determines metabolic health effects in mice. Cell metabolism 33, 1808–1819.

Manning, R. (2001). Fatty acids in pollen: a review of their importance for honey bees. Bee world 82, 60–75.

Moriyama, M. and Fukatsu, T. (2022). Host’s demand for essential amino acids is compensated by an extracellular bacterial symbiont in a hemipteran insect model. Frontiers in Physiology 13, 1028409.

Murray, E. A., Bossert, S. and Danforth, B. N. (2018). Pollinivory and the diversification dynamics of bees. Biology letters 14, 20180530.

Nicolson, S. W. (2022). Sweet solutions: nectar chemistry and quality. Philosophical Transactions of the Royal Society B 377, 20210163.

Paoli, P. P., Donley, D., Stabler, D., Saseendranath, A., Nicolson, S. W., Simpson, S. J. and Wright, G. A. (2014a). Nutritional balance of essential amino acids and carbohydrates of the adult worker honeybee depends on age. Amino acids 46, 1449–1458.

Paoli, P. P., Wakeling, L. A., Wright, G. A. and Ford, D. (2014b). The dietary proportion of essential amino acids and Sir2 influence lifespan in the honeybee. Age 36, 1239–1247.

Pedroso, J. A., Zampieri, T. T. and Donato Jr, J. (2015). Reviewing the effects of L-leucine supplementation in the regulation of food intake, energy balance, and glucose homeostasis. Nutrients 7, 3914–3937.

Peters, B., Keller, A. and Leonhardt, S. D. (2022). Diets maintained in a changing world: Does land-use intensification alter wild bee communities by selecting for flexible generalists? Ecology and Evolution 12, e8919.

Piper, M. D., Soultoukis, G. A., Blanc, E., Mesaros, A., Herbert, S. L., Juricic, P., He, X., Atanassov, I., Salmonowicz, H. and Yang, M. (2017). Matching dietary amino acid balance to the in silico-translated exome optimizes growth and reproduction without cost to lifespan. Cell metabolism 25, 610–621.

Prosser, C. L. (1989). Two pathways of evolution of neurotransmitters-modulators. In Evolution of the first nervous systems, pp. 177–193. Springer.

Prosser, W. and Douglas, A. (1991). The aposymbiotic aphid: an analysis of chlortetracycline-treated pea aphid, Acyrthosiphon pisum. Journal of Insect Physiology 37, 713–719.

Raubenheimer, D. and Simpson, S. J. (1999). Integrating nutrition: a geometrical approach.pp. 67–82. Springer.

Reifenberg, P. and Zimmer, A. (2024). Branched-chain amino acids: physico-chemical properties, industrial synthesis and role in signaling, metabolism and energy production. Amino Acids 56, 51.

Roulston, T. H. and Cane, J. H. (2000). Pollen nutritional content and digestibility for animals. Plant systematics and Evolution 222, 187–209.

Sereia, M. J., Toledo, V. de A. A. de, Furlan, A. C., Faquinello, P., Maia, F. M. C. and Wielewski, P. (2013). Alternative sources of supplements for Africanized honeybees submitted to royal jelly production. Acta Scientiarum. Animal Sciences 35, 165–171.

Sheiner, J. B., Morris, P. and Anderson, G. H. (1985). Food intake suppression by histidine. Pharmacology Biochemistry and Behavior 23, 721–726.

Simpson, S. J. and Raubenheimer, D. (2005). Obesity: the protein leverage hypothesis. obesity reviews 6, 133–142.

Somerville, D. and Nicol, H. (2006). Crude protein and amino acid composition of honey bee-collected pollen pellets from south-east Australia and a note on laboratory disparity. Australian Journal of Experimental Agriculture 46, 141–149.

Stabler, D., Paoli, P. P., Nicolson, S. W. and Wright, G. A. (2015). Nutrient balancing of the adult worker bumblebee (Bombus terrestris) depends on the dietary source of essential amino acids. The Journal of experimental biology 218, 793–802.

Stabler, D., Power, E. F., Borland, A. M., Barnes, J. D. and Wright, G. A. (2018). A method for analysing small samples of floral pollen for free and protein-bound amino acids. Methods in Ecology and Evolution 9, 430–438.

Stabler, D., Al-Esawy, M., Chennells, J. A., Perri, G., Robinson, A. and Wright, G. A. (2021). Regulation of dietary intake of protein and lipid by nurse-age adult worker honeybees. Journal of Experimental Biology 224, jeb230615.

Vaudo, A. D., Stabler, D., Patch, H., Tooker, J., Grozinger, C. and Wright, G. (2016). Bumble bees regulate their intake of essential protein and lipid pollen macronutrients. Journal of Experimental Biology 219, 3962–3970.

Vaudo, A. D., Tooker, J. F., Patch, H. M., Biddinger, D. J., Coccia, M., Crone, M. K., Fiely, M., Francis, J. S., Hines, H. M. and Hodges, M. (2020). Pollen protein: lipid macronutrient ratios may guide broad patterns of bee species floral preferences. Insects 11, 132.

Weiner, C. N., Hilpert, A., Werner, M., Linsenmair, K. E. and Blüthgen, N. (2010). Pollen amino acids and flower specialisation in solitary bees. Apidologie 41, 476–487.

Wu, T., Baatar, D., O’Connor, A. E., O’Bryan, M. K., Stringer, J. M., Hutt, K. J., Aponso, M. M., Monro, K., Luo, J. and Zhu, Y. (2024). Exome-informed formulations of food proteins enhance body growth and feed conversion efficiency in ad libitum-fed mice. Food Research International 176, 113819.

Yokota, S. C., Broeckling, C. and Hs, A. (2024). Pollen foraging preferences in honey bees and the nutrient profiles of the pollen. Scientific Reports 14, 15028.

