## Supplement tables and figures for "Nutrition of honeybees is constrained by the ratios of essential amino acids in pollen protein"

**Supplementary tables and figures**

**Table S1.** Canonical discriminant analysis of essential amino acids quantified in bee tissues, stored pollen (bee bread), and hand collected, fresh pollen.

| | Essential amino acids | | | |
|---|---|---|---|---|
| | Canonical discriminant function statistics | | | |
| Function | Eigenvalue | % variance | test stat | P value |
| 1 | 5881 | 95.9 | $x_{14}^2$=56.85 | <0.001 |
| 2 | 252 | 4.1 | $x_6^2$=22.13 | 0.001 |

Pooled within-groups correlations

| | Function 1 | Function 2 |
|---|---|---|
| Valine | 0.185* | -0.017 |
| Leucine | 0.082* | -0.012 |
| Phenylalanine | -0.031* | -0.025 |
| Isoleucine | 0.027* | -0.026 |
| Arginine | -0.022* | -0.005 |
| Histidine | -0.006* | 0.001 |
| Threonine | 0.021 | 0.302* |
| Methionine | -0.002 | -0.033* |
| Lysine | 0.00 | -0.022* |

Canonical discriminant function coefficients

| | Function 1 | Function 2 |
|---|---|---|
| Bee tissue | 76.9 | -3.30 |
| Stored pollen | -13.7 | 39.7 |
| Fresh pollen | -58.8 | -5.30 |

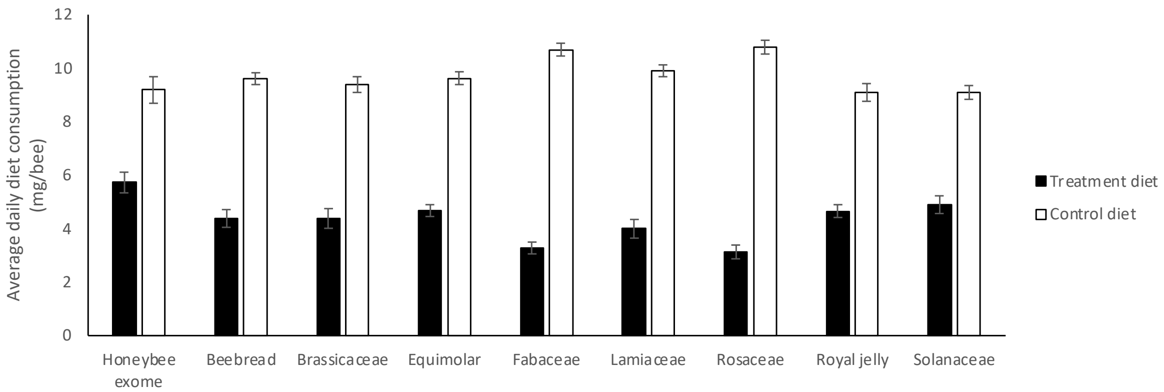


**Figure S1.** Average daily consumption of treatment and control diets from feeding experiments. Treatment diets contained 3% free essential amino acids matching the profile of the treatment name. Control diets contained all other nutrients as the treatment diet, excluding free amino acids. The nutrient space was backfilled with maltodextrin. Error bars represent standard error of mean. N = 20 cohorts of 30 bees per treatment.

**Table S2.** Measured essential and non-essential amino acids from bee, royal jelly, and pollen tissues.

| | EAA (µg/mg) | | | | | | | | | | Non-EAA (µg/mg) | | | | | | | | |
|---|---|---|---|---|---|---|---|---|---|---|---|---|---|---|---|---|---|---|---|
| **Tissue type** | **arg** | **his** | **ile** | **leu** | **lys** | **met** | **phe** | **thr** | **trp** | **val** | **ala** | **asp** | **cys** | **glu** | **gln** | **pro** | **ser** | **tyr** | **Total (µg/mg)** |
| Honeybee pupae | 19.20 | 23.60 | 20.80 | 32.40 | 27.20 | 8.17 | 14.10 | 16.40 | 0.00 | 24.00 | 26.40 | 33.80 | 4.50 | 50.80 | 23.10 | 27.90 | 18.40 | 21.10 | 391.87 |
| Royal jelly | 17.30 | 12.80 | 17.70 | 27.20 | 23.00 | 4.98 | 15.30 | 15.80 | 0.00 | 20.20 | 10.40 | 61.10 | 2.44 | 34.50 | 10.20 | 15.30 | 20.60 | 9.21 | 318.03 |
| Brassicaceae | 42.05 | 4.60 | 1.92 | 8.42 | 28.42 | 12.01 | 3.90 | 3.15 | 0.00 | 8.42 | 14.63 | 1.90 | 12.11 | 30.80 | 0.01 | 10.02 | 19.97 | 13.45 | 215.78 |
| Fabaceae | 28.01 | 25.80 | 3.04 | 5.66 | 5.90 | 2.79 | 7.01 | 2.27 | 0.00 | 2.34 | 11.78 | 7.64 | 11.29 | 15.68 | 0.00 | 5.32 | 12.24 | 5.57 | 152.33 |
| Lamiaceae | 18.38 | 22.37 | 0.12 | 5.32 | 5.61 | 3.12 | 18.34 | 1.81 | 0.00 | 2.64 | 2.33 | 10.77 | 7.68 | 12.97 | 0.00 | 4.23 | 12.51 | 3.72 | 131.91 |
| Rosaceae | 38.48 | 35.89 | 1.52 | 5.87 | 12.06 | 3.61 | 3.76 | 2.66 | 0.00 | 4.57 | 6.76 | 6.84 | 9.77 | 23.12 | 0.00 | 8.17 | 12.80 | 5.85 | 181.76 |
| Solanaceae | 11.39 | 0.94 | 0.51 | 5.03 | 14.18 | 8.48 | 17.19 | 1.81 | 0.00 | 1.38 | 2.48 | 0.69 | 7.38 | 12.28 | 0.00 | 3.67 | 11.19 | 3.78 | 102.37 |

**Table S3** Proportions (%) of essential amino acids in each treatment group.

| | Equal EAA | Honeybee exome | Royal jelly | Bee bread | Brassicaceae | Fabaceae | Lamiaceae | Rosaceae | Solanaceae |
|---|---|---|---|---|---|---|---|---|---|
| arg | 10.0 | 10.9 | 10.7 | 24.1 | 36.1 | 32.8 | 22.9 | 34.4 | 18.1 |
| his | 10.0 | 5.0 | 7.9 | 16.6 | 3.9 | 30.2 | 27.9 | 32.1 | 1.5 |
| ile | 10.0 | 13.6 | 10.9 | 1.6 | 1.6 | 3.6 | 0.1 | 1.4 | 0.8 |
| leu | 10.0 | 18.4 | 16.8 | 9.7 | 7.2 | 6.6 | 6.6 | 5.3 | 8.0 |
| lys | 10.0 | 14.1 | 14.2 | 8.0 | 24.4 | 6.9 | 7.0 | 10.8 | 22.6 |
| met | 10.0 | 4.7 | 3.1 | 0.5 | 10.3 | 3.3 | 3.9 | 3.2 | 13.5 |
| phe | 10.0 | 7.5 | 9.5 | 4.7 | 3.3 | 8.2 | 22.9 | 3.4 | 27.4 |
| thr | 10.0 | 12.1 | 9.8 | 29.2 | 2.7 | 2.7 | 2.3 | 2.4 | 2.9 |
| val | 10.0 | 11.6 | 12.5 | 2.6 | 7.2 | 2.7 | 3.3 | 4.1 | 2.2 |
| trp | 10.0 | 2.2 | 4.8 | 3.1 | 3.1 | 3.1 | 3.1 | 3.1 | 3.1 |
| Total | 100 | 100 | 100 | 100 | 100 | 100 | 100 | 100 | 100 |

**Table S4**. Proportions (%) of ingredients used to make treatment and control diets for all feeding experiments.

| Diet component | EAA treatment diets | Control diets |
|---|---|---|
| Powdered sugar | 20.25 | 20.25 |
| Invert syrup | 60.62 | 60.62 |
| Free EAAs | 3 | 0 |
| Emulsion | 4.11 | 4.11 |
| Maltodextrin | 12.03 | 15.03 |
| Total composition (%) | 100.0 | 100.0 |

**Table S5.** Essential amino acid proportions (%) of three representative profiles of honeybee tissue, the honeybee exome, the proportion of essential amino acids estimated by de Groot, 1953, and those rendered from honeybee pupae.

| | *A. mel* Exome | de Groot | Worker pupae |
|---|---|---|---|
| arg | 10.9 | 10.7 | 10.2 |
| his | 5.0 | 14.3 | 10.3 |
| ile | 13.6 | 5.4 | 10.7 |
| leu | 18.4 | 16.1 | 16.8 |
| lys | 14.1 | 14.3 | 13.9 |
| met | 4.7 | 8.9 | 4.4 |
| phe | 7.5 | 10.7 | 8.4 |
| thr | 12.1 | 5.4 | 8.3 |
| val | 11.6 | 10.7 | 12.1 |
| trp | 2.2 | 3.6 | 4.8 |
| Total | 100 | 100 | 100 |

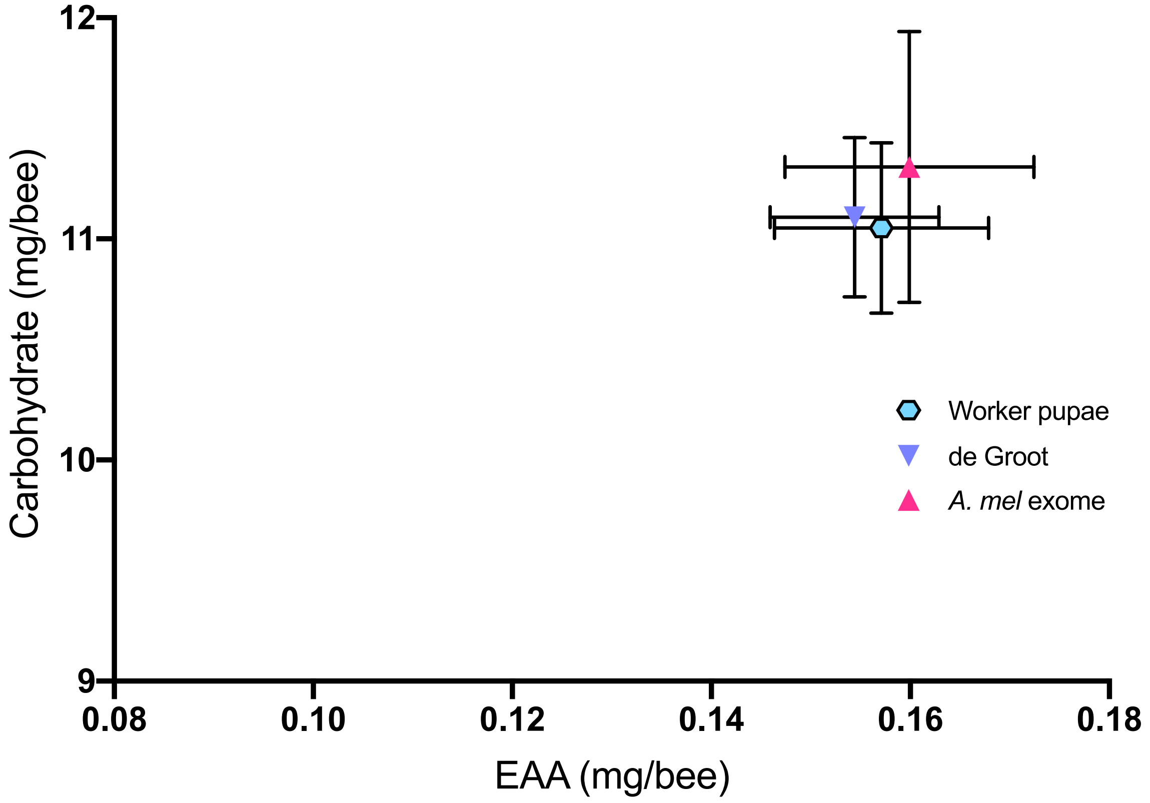


**Figure S2.** Essential amino acid (EAA) vs carbohydrate consumption of diets reflecting three profiles of honeybee EAA composition. Error bars represent standard error of mean. N = 10 cohorts of 30 bees per treatment.

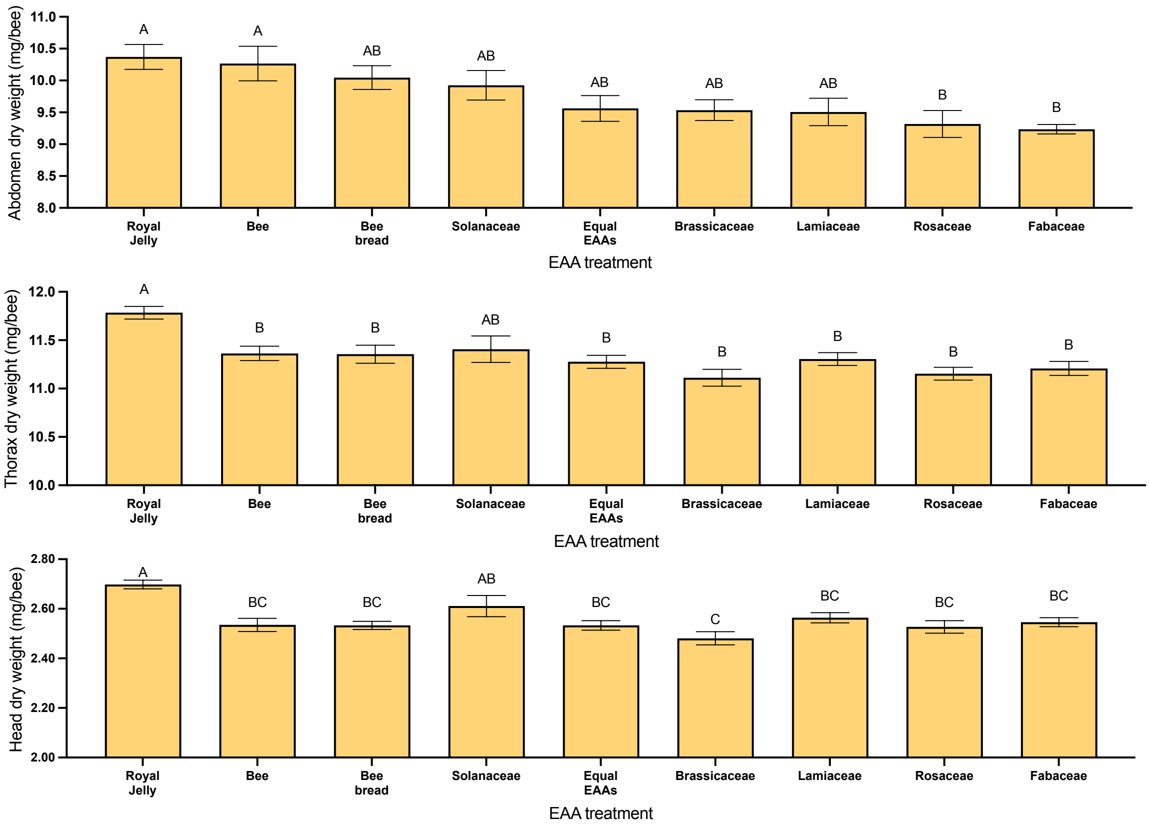


**Figure S3.** Dry weights of segmented bee abdomens, thoraxes, and heads from feeding experiments where treatment diets reflected essential amino acid (EAA) profiles of bee, royal jelly, pollen tissues, or an equal proportion of EAAs. Error bars represent standard error of mean. N = 10 cohorts of 10 abdomens, thoraxes, and heads per treatment. Bees fed diets based on the EAA composition of royal jelly and bee tissues had abdomens 1.11 times heavier than bees on the diets of rosacea and fabaceae EAAs (Šidák's *post hoc*, $P's \leq 0.045$). Bee thorax weight was largely unaffected by the EAA composition of diet, though the thorax of bees feeding on royal jelly EAAs were ~1.05 times heavier than all other treatments (Šidák's *post hoc*, $P's \leq 0.023$), except Solanaceae pollen EAAs, which were similar to bees fed royal jelly EAAs (Šidák's *post hoc*, $P = 0.072$). Bees fed royal jelly EAAs also had heads that weighed ~1.07 times more than all other treatments (Šidák's *post hoc*, $P's \leq 0.002$), except for solanaceae, which weighed similarly to royal jelly (Šidák's *post hoc*, $P = 0.431$) and weighed more than the heads of bees fed EAAs of brassicaceae pollen (Šidák's *post hoc*, $P = 0.014$).

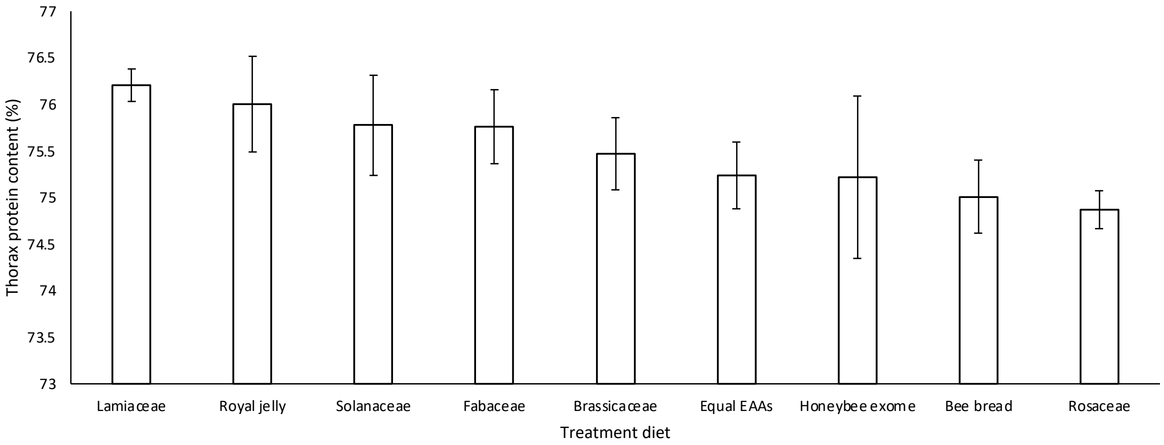


**Figure S4.** Average protein content of thoraxes from bees fed treatment diets. Error bars represent standard error of mean. N = 10 cohorts of 10 thoraxes per treatment.

**Table S6.** Proportion of EAAs in formulated diets representing their relative loadings from a PCA of pollen, bee bread, and honeybee tissues and products. Values in green sections highlight the amino acids that were manipulated in each experiment.

| **EAAs (%)** | **Factor 1A** | **Factor 1B** | **Factor 2A** | **Factor 2B** | **Factor 3A** | **Factor 3B** |
|---|---|---|---|---|---|---|
| Arginine | 1 | 9 | 11 | 11 | 9 | 11 |
| Histidine | 1 | 9 | 11 | 11 | 9 | 11 |
| Isoleucine | 6 | 1 | 11 | 11 | 9 | 11 |
| Leucine | 6 | 1 | 11 | 11 | 9 | 11 |
| Lysine | 16 | 16 | 9 | 1 | 9 | 11 |
| Methionine | 16 | 16 | 9 | 1 | 9 | 11 |
| Phenylalanine | 16 | 16 | 11 | 11 | 18 | 2 |
| Threonine | 16 | 16 | 2 | 18 | 9 | 11 |
| Valine | 6 | 1 | 11 | 11 | 9 | 11 |
| Tryptophan | 16 | 16 | 11 | 11 | 9 | 11 |
| Total | 100 | 100 | 100 | 100 | 100 | 100 |

**Table S7.** Proportions of essential amino acids (EAAs) in diets for each treatment group, where eight EAAs were in equal proportion and the proportion of two EAAs varied in each treatment. Values in green sections highlight the amino acids manipulated in each experiment.

| % of EAA composition | Control | His:Leu | | | | His:Val | | | | His:Ile | | | | His:Lys | | | | Leu:Lys | | | | Leu:Arg | | | |
|---|---|---|---|---|---|---|---|---|---|---|---|---|---|---|---|---|---|---|---|---|---|---|---|---|---|
| | (equal EAAs) | 18:2 | 14:6 | 6:14 | 2:18 | 18:2 | 14:6 | 6:14 | 2:18 | 18:2 | 14:6 | 6:14 | 2:18 | 18:2 | 14:6 | 6:14 | 2:18 | 18:2 | 14:6 | 6:14 | 2:18 | 18:2 | 14:6 | 6:14 | 2:18 |
| Arginine | 10 | 10 | 10 | 10 | 10 | 10 | 10 | 10 | 10 | 10 | 10 | 10 | 10 | 10 | 10 | 10 | 10 | 10 | 10 | 10 | 10 | 2 | 6 | 14 | 18 |
| Histidine | 10 | 18 | 14 | 6 | 2 | 18 | 14 | 6 | 2 | 18 | 14 | 6 | 2 | 18 | 14 | 6 | 2 | 10 | 10 | 10 | 10 | 10 | 10 | 10 | 10 |
| Iso-leucine | 10 | 10 | 10 | 10 | 10 | 10 | 10 | 10 | 10 | 2 | 6 | 14 | 18 | 10 | 10 | 10 | 10 | 10 | 10 | 10 | 10 | 10 | 10 | 10 | 10 |
| Leucine | 10 | 2 | 6 | 14 | 18 | 10 | 10 | 10 | 10 | 10 | 10 | 10 | 10 | 10 | 10 | 10 | 10 | 18 | 14 | 6 | 2 | 18 | 14 | 6 | 2 |
| Lysine | 10 | 10 | 10 | 10 | 10 | 10 | 10 | 10 | 10 | 10 | 10 | 10 | 10 | 2 | 6 | 14 | 18 | 2 | 6 | 14 | 18 | 10 | 10 | 10 | 10 |
| Methionine | 10 | 10 | 10 | 10 | 10 | 10 | 10 | 10 | 10 | 10 | 10 | 10 | 10 | 10 | 10 | 10 | 10 | 10 | 10 | 10 | 10 | 10 | 10 | 10 | 10 |
| Phenylalanine | 10 | 10 | 10 | 10 | 10 | 10 | 10 | 10 | 10 | 10 | 10 | 10 | 10 | 10 | 10 | 10 | 10 | 10 | 10 | 10 | 10 | 10 | 10 | 10 | 10 |
| Threonine | 10 | 10 | 10 | 10 | 10 | 10 | 10 | 10 | 10 | 10 | 10 | 10 | 10 | 10 | 10 | 10 | 10 | 10 | 10 | 10 | 10 | 10 | 10 | 10 | 10 |
| Tryptophan | 10 | 10 | 10 | 10 | 10 | 10 | 10 | 10 | 10 | 10 | 10 | 10 | 10 | 10 | 10 | 10 | 10 | 10 | 10 | 10 | 10 | 10 | 10 | 10 | 10 |
| Valine | 10 | 10 | 10 | 10 | 10 | 2 | 6 | 14 | 18 | 10 | 10 | 10 | 10 | 10 | 10 | 10 | 10 | 10 | 10 | 10 | 10 | 10 | 10 | 10 | 10 |
| Total (%) | 100 | 100 | 100 | 100 | 100 | 100 | 100 | 100 | 100 | 100 | 100 | 100 | 100 | 100 | 100 | 100 | 100 | 100 | 100 | 100 | 100 | 100 | 100 | 100 | 100 |

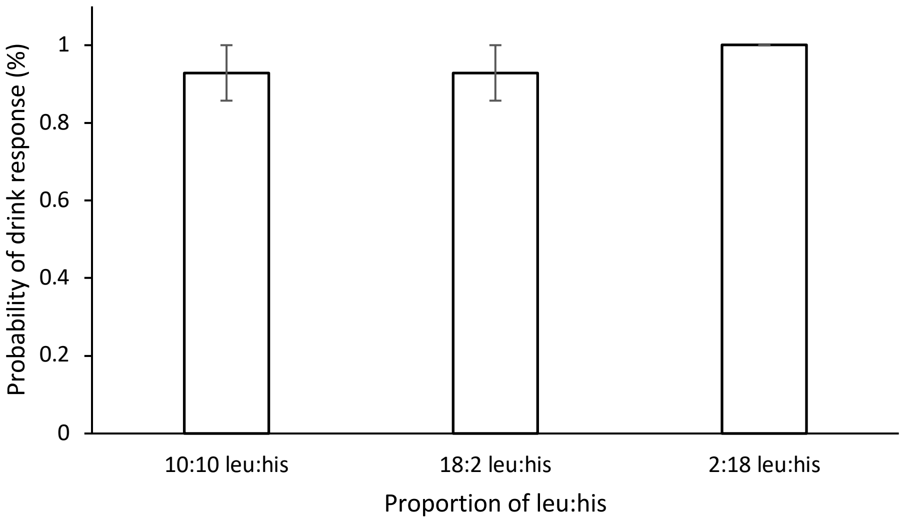


**Figure S5.** Probability of drink response for bees presented solutions varying in proportions of leucine and histidine. Error bars represent standard error of mean. N = 14 bees per treatment.
